# Urbanisation impacts the diversity, coloration, and body size of wild bees in a Mediterranean city

**DOI:** 10.1101/2022.12.09.519739

**Authors:** Arnaud Badiane, Lise Ropars, Floriane Flacher, Lucie Schurr, Marie Zakardjian, Laurence Affre, Magali Deschamps-Cottin, Sophie Gachet, Christine Robles, Benoît Geslin

## Abstract

Urbanisation is a growing phenomenon causing the decline of wild bees globally. Yet, bees manage to persist in the urban matrix thanks to islands of vegetation in public parks and private gardens. While we begin to comprehend the impact of urbanisation on bees’ diversity and abundance, our understanding of its impact on the functional diversity of wild bees is limited. Here, we use an integrative approach to investigate the response of wild bees to urbanisation at the community, species, and individual levels. To do so, we sampled wild bees in 24 public parks along an urbanisation gradient in the Mediterranean city of Marseille. We found that species richness and abundance decreased in more urbanised areas, but increased in larger city parks. Moreover, larger individuals within species, but not larger species, were found in larger city parks, suggesting that park size is crucial for the persistence of bees in cities. Interestingly, we show that brighter species were found in parks surrounded by a large amount of impervious surface, highlighting the importance of colour traits in the response to environmental changes. Finally, our results revealed that larger species, but not larger individuals, were also more colourful. In summary, our study not only confirmed that urbanisation negatively impacts community-level traits, but that it also affects species’ coloration and individuals’ body size, thus improving our understanding of the functional response of wild bees to urbanisation. We suggest that increasing park size may compensate for the negative effects of urbanisation on wild bees.

## Introduction

Bees constitute one of the major groups of pollinators of wild plants and crops worldwide (Potts et al. 2016; Hung et al. 2018; Zattara and Aizen 2021). Over the past sixty years, bees experienced a sharp decline globally (Zattara and Aizen 2021). Several anthropogenic factors are responsible for this decline, including urbanisation and agricultural intensification causing habitat and floral resource loss, the use of pesticides, parasites, the introduction of invasive species and climate change (Potts et al. 2010; Goulson et al. 2015; Sánchez-Bayo and Wyckhuys 2019). Among these causes, urbanisation is especially preoccupying because urban areas are growing at an unprecedented rate (United Nations, 2018), transforming semi-natural and agricultural habitats into impervious surfaces (McKinney 2002) detrimental to bees (Cardoso and Gonçalves 2018; Baldock 2020). Yet, islands of vegetation subsist in urban landscapes, such as private gardens, allotments, and public parks, allowing bees to persist in these environments (Baldock et al. 2015, 2019; Geslin et al. 2015; Theodorou et al. 2020). The urban matrix therefore acts as an environmental filter and its permeability, which can be highly variable among cities, depends on the amount, extent, quality, and degree of isolation of these islands of vegetation (Mcintyre and Hostetler 2001; Braaker et al. 2014; Fattorini 2016; Banaszak-Cibicka et al. 2018). When city parks are managed so as to offer favourable conditions for bee assemblages, urban environments can harbour a bee species diversity and abundance comparable to what is found in natural habitats, but not necessarily in terms of functional diversity (Hall et al. 2017; Banaszak-Cibicka et al. 2018). An increasing amount of work focuses on the functional aspects of urban impacts on bees, examining the morphological and life-history traits allowing or preventing bees to cope with urban environments (e.g. Geslin et al. 2013, 2016; Zaninotto et al. 2021) and references therein).

Several functional traits have been found to promote the presence of bees in large cities. Indeed, social behaviour, broad dietary niche (i.e., polylectism), cavity-nesting habits, and early spring phenology seem to be favoured in urban landscapes whereas solitary and parasitic behaviours, narrow dietary niche (i.e., oligolectism), ground-nesting habits, and late spring phenology appear to be unsuccessful traits in cities (Zanette et al. 2005; Hernandez et al. 2009; Banaszak-Cibicka et al. 2018; Buchholz et al. 2020; Ayers and Rehan 2021; Zaninotto et al. 2021). Regarding body size, however, evidence is more contrasted. On one hand, some studies found that large-sized species decreased in abundance and diversity in urban environments (Banaszak-Cibicka and Żmihorski 2012; Geslin et al. 2016; Banaszak-Cibicka et al. 2018) possibly because large body size correlates with extinction risk in insects (e.g., Nolte et al. 2019). On the other hand, other studies found that small species were less common in urban centres because of their reduced dispersal abilities whereas large-sized species were less affected by urbanisation as they have good flight abilities allowing them to penetrate the urban matrix and hop from a suitable patch to another (Gathmann and Tscharntke 2002; Ahrné et al. 2009). Reduced flight abilities generally make smaller bees less mobile and more sensitive to habitat fragmentation in general than larger bee species (e.g., Steffan-Dewenter and Tscharntke 1999; Greenleaf et al. 2007; Warzecha et al. 2016; Gérard et al. 2021). Interestingly, body size correlates with several phenotypic traits fulfilling important ecological functions. For instance, small-sized bees usually have small mouthparts, which associates with a narrower dietary niche because they cannot exploit some types of flowers (e.g., tubular - Stang et al. 2006) whereas the reverse is true for larger species. It is therefore pertinent to use body size when assessing the effect of urbanisation on the functional diversity of bees (Theodorou et al. 2021).

Urbanisation may also affect other traits that play important functions in bees, such as coloration. Bees indeed display a great variety of colours. Some species are entirely black or darkly coloured, while others display bright colours including yellow, orange, red, green, blue, violet, and white (Michez et al. 2019). These colour traits play various functions. Bright colours often act as Mullerian and Batesian aposematic signals in bees (Badejo et al. 2020), especially when black coloration associates with bright stripes (Mappes et al. 2005; Caro and Ruxton 2019). Melanin pigments responsible for the dark coloration can also contribute to defence functions by encapsulating pathogens (Siva-Jothy et al. 2005) and protecting against UV radiations (Badejo et al. 2020). Body coloration can also serve as camouflage (Williams 2007) and play a role in thermoregulation processes, for example via the thermal melanism hypothesis (Clusella Trullas et al. 2007) stating that darker colours should be favoured in colder environments. Hence, given the functional importance of body coloration in bees, urbanisation can affect bee coloration via its impact on the multiple processes involving colour traits. For instance, urbanisation reduces predation pressures (Lagucki et al. 2017; Eötvös et al. 2018, 2020), which in turn may affect aposematic signals (Valkonen et al. 2012). Moreover, the urban heat-island effect in cities (Memon et al. 2008) impacts water balance and thermoregulation processes of bees (Hamblin et al. 2017) such that bee species are close to their critical thermal limit and/or their critical water content (Burdine and McCluney 2019a). Thus, we could hypothesise that darker species reach their critical thermal limit faster than brighter ones, which would make them less successful in cities, especially in cities located in warm regions. Finally, urban landscapes have a different background colour than surrounding natural habitats due to buildings and impervious surfaces, and this may alter camouflage and colour signal efficacy (Delhey and Peters 2017). Because the selective forces affecting coloration detailed above have conflicting effects, it is challenging to predict how urbanisation will affect bee coloration. Thus, exploring whether urban environments promote or hinder colourful traits in bees will bring new insights into the ecological impacts of urbanisation processes.

In this study, we aimed to assess the impact of urbanisation on wild bee assemblages in the Mediterranean city of Marseille, France. While the Mediterranean region is a hotspot for bee diversity (Nielsen et al. 2011; Ropars et al. 2020a), it also suffers from anthropogenic pressures including increasing urbanisation (García-Nieto et al. 2018), enhancing the need to improve our understanding of the response of bees to urbanisation. Our study focuses on wild bees only and excludes the honey bee (*Apis mellifera*) because the latter is a non-native, managed species with possible negative impacts on wild bee communities (Ropars et al. 2019, 2020b). We sampled urban parks along an urbanisation gradient in Marseille in order to investigate the extent to which landscape variables related to urbanisation affect wild bees at the community level (i.e. species diversity and abundance), at the species level (i.e. mean specific body size and coloration), and at the individual level (i.e. within species variation in body size and coloration). This study design allows us to assess the impact of urbanisation at three biological scales so that we can improve our understanding of the response of wild bees, and other species, to urbanisation. In addition, we also explored the relationship between body size and coloration in wild bees, both at the inter- and intra-specific level, as it has never been empirically studied in the past.

## Material and Methods

### Study sites

The study was conducted in the Mediterranean city of Marseille (France) during the spring and summer of 2016 (April to July) for fieldwork and during the spring of 2020 for laboratory analyses. With 240 km^2^ and 871,103 inhabitants (INSEE, 2020), Marseille is the second-largest and one of the oldest cities of France. The region is characterised by a Mediterranean climate with cool winters and hot summers accompanied by irregular precipitations in spring and autumn and pronounced summer drought. In contrast to most European cities, Marseille is not surrounded by agricultural crops but by calcareous massifs dominated by biodiversity-rich areas such as shrublands. This configuration thus offers interesting gradients from natural habitats to highly urbanised areas (Lizée et al. 2012; Lizee et al. 2016), which is ideal to study how animals cope with urbanisation.

We selected 22 city parks and 2 university campus (similarly managed) covering an urbanisation gradient within the city of Marseille, from the highly urbanised city centre to less urbanised areas on the periphery (Figure 1). These parks vary in size (range 1-31 ha, mean = 9 ha), and offer various land-use contexts, with various amounts of surrounding vegetation and impervious surface, and various degrees of isolation from natural areas, as the distance from a park to the closest natural areas ranges from 0.5 km to 7.5 km. One urban park was excluded from the following analyses because no native bees were found foraging in the park (only *Apis mellifera*).

**Figure 1.**
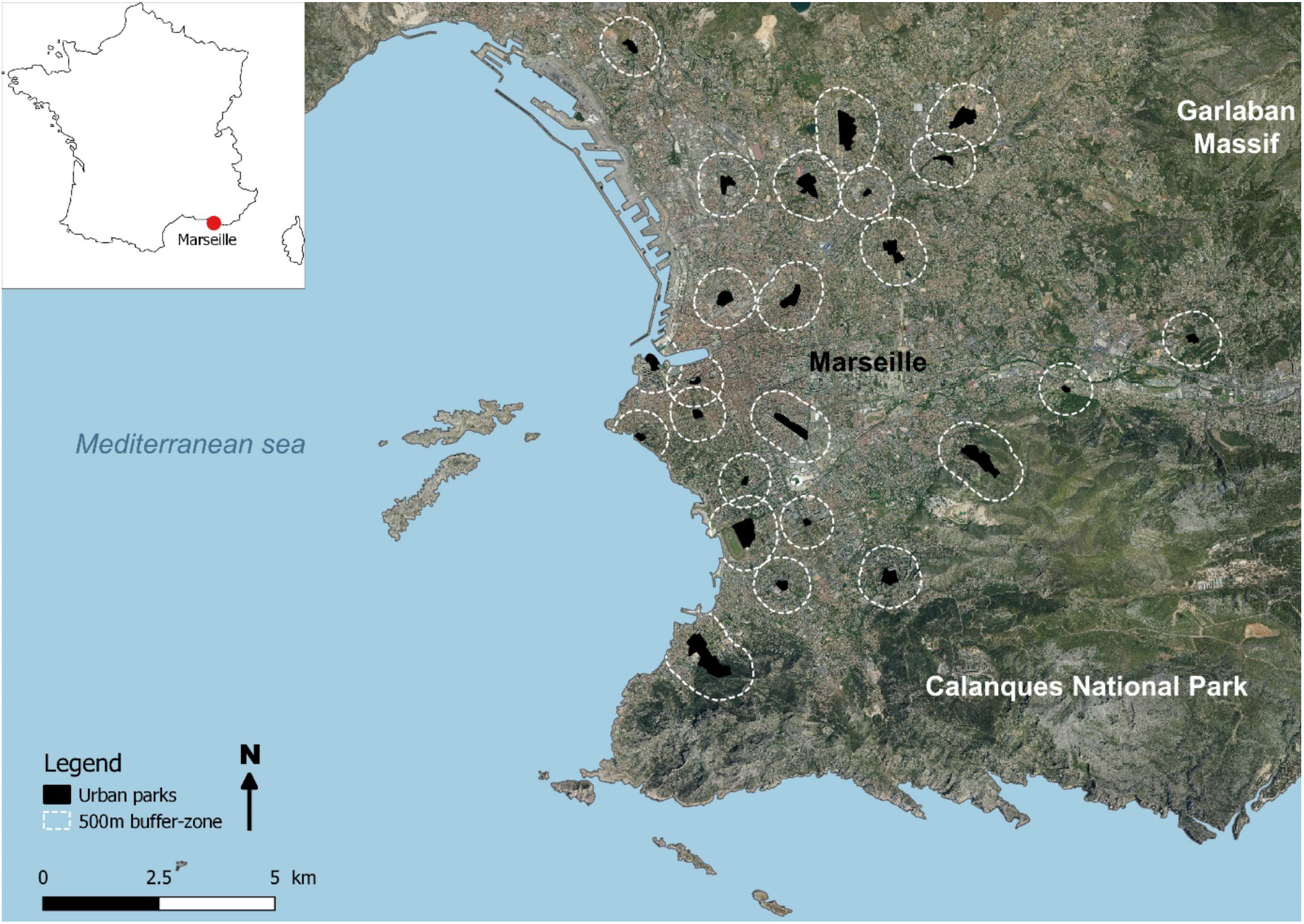
Location of the 24 urban parks (in black) sampled within the city of Marseille with their 500m buffer-zone (white dotted lines).

### Landscape variables

To characterize landscape variables, we used the land cover map from Lizée et al. (2012) built with SPOT and IGN data (SPOT5 - 2004; BD1000-2006; BD Carto® IGN - 2004). We combined these data using QGIS software on a 10 m-resolution raster map and contains 5 classes: impervious surface, rocky habitat, sparsely vegetated area, herbaceous stratum, tree stratum.

First, we calculated the distance from each park to the closest natural area by drawing a straight line between the park and the closest natural area, which in Marseille corresponds mostly to the closest mountain range. Then, we created a polygon around each of the 24 parks and calculated their area. We also drew a buffer-zone of 500 m around each of the 24 urban parks in order to calculate the area of each land cover class, and made sure to exclude the area within the parks. We chose a 500-m buffer because it encompasses the mean foraging range of most wild bee species we observed as mentioned in Wright et al. (2015). We counted the number of pixels of each class in the 500-m band around each urban park. Then, we combined rocky habitats and impervious surfaces as one class and three vegetation classes (i.e., grasses, scarce vegetations, and trees) all together to only have two classes: impervious surfaces and vegetation surfaces.

### Bee sampling and pollination network description

In each of the 24 parks, we surveyed 16 transects of 10 m during five minutes at a pace of one meter every 30 seconds. To maximize the bee species richness, we placed eight transects along a linear of shrub or bush, and eight transects within a lawn totalizing 384 transects. We prospected each transect three times during the period April to July 2016 for a total of 1152 transect visits.

We captured with a net all wild bees observed foraging within 2 meters on both sides of each transect. We identified each plant species on which bees were foraging. Bee specimens collected were pinned and dried prior to identification by professional taxonomists (E. Dufrêne for cuckoo bee species, D. Genoud for Andrenidae, Anthophorinii, *Colletes* sp. and Halictidae and M. Aubert for Megachilidae, Ceratinii and *Hylaeus sp*.).

To evaluate the completeness of our samplings and estimate the potential maximum bee species richness in the city of Marseille, we used the Chao and jackknife indexes including captures and observations on plant species (Gotelli and Colwell 2011). We calculated these indexes using the function *ChaoSpecies* within the *Spade-R* package in R version 3.6 software (Chao et al. 2016).

### Body size and colour variables

We took calibrated photographs of the dorsal part of each captured bee using a DSLR Nikon D500 mounted with a Tokina 100-mm macro lens. For each photograph, we placed a millimetric scale and a colour chart with a grey scale (i.e., SpyderCheckr, Datacolor Inc.). Then, we imported the pictures in raw format in the software ImageJ (Schneider et al. 2012) and used the ‘line’ tool to measure the intertegular span of each individual, which is a reliable proxy of body size in bees (Cane 1987).

To objectively assess bee coloration, we used the Quantitative Colour Pattern Analysis (QCPA) framework (van den Berg et al. 2020) implemented in the Multispectral Image Analysis and Calibration (MICA) Toolbox (Troscianko and Stevens 2015), an ImageJ plugin. First of all, we created a cone-catch model for our camera setup using a colour chart (X-Rite colorCheckr passport) of known reflectance. This step allows us to convert the RGB values recorded by our photography setup into the standardised colorimetric values of the CIELAB colour space. We used the CIELAB, a colour space based on human vision, because we did not have access to the UV range, and since bees are capable of UV vision, we could not use the bee visual system (Menzel and Blakers 1976; Peitsch et al. 1992). CIELAB is a three-dimensional colour space in which each colour is defined by three chromatic variables or coordinates: L*, a* and b*. Lightness (L*) is the percentage of light reflected from a surface and goes from black (0) to white (100). The coordinate a* corresponds to a green-to-red colour variation and coordinate b* corresponds to a blue-to-yellow colour variation. Then, we generated a multispectral image from RAW photographs using the MICA Toolbox and adjusted the white balance with the 96% white standard from the colour chart. We then selected two body regions of interest to be measured, namely the thorax and the abdomen, by surrounding these body parts, excluding wings and artefacts such as the entomological pin. After having converted our multispectral image into the CIELAB cone-catch model, we obtained the mean L*a*b* values for the whole thorax and the whole abdomen of each individual. Finally, we calculated the L*a*b* values of the entire body by taking the averaged values between the thorax and the abdomen, therefore characterising the body coloration of each individual.

### Statistical analyses

To better understand the impact of urbanisation on wild bee communities, we explored the relationships between urbanisation variables, community-level variables (i.e., species diversity, abundance) and individual-level variables (i.e., body size, coloration) using a piecewise structural equation modelling (SEM) in combination with (generalised) linear (mixed-effects) models. To do so, we used R v.3.6.2 (R Core Team 2019) with the R packages *piecewiseSEM* v2.1 package (Lefcheck 2016) and *nlme* (Pinheiro et al. 2019). SEM is a suitable tool to evaluate direct and indirect effects in descriptive analyses of ecological systems (Grace et al. 2010). In addition, piecewise SEM tests for missing paths between variables using Shipley’s test of d-separation (Shipley 2013), allowing us to adjust our initial model to improve its fit and biological significance. Adequate model goodness-of-fit is first indicated by a non-significant p-value based on the Chi-squared test (Shipley 2009). Then, goodness-of-fit can be improved using a combination of indices, including Akaike’s Information Criterion corrected for small sample size (AICc) obtained from Fisher’s C statistic, and the Bayes-Schwarz Information Criterion (BIC), the latter being the most reliable for model selection using piecewise SEM (Hertzog 2018).

We built two similar models that differ in the way individual-level variables are accounted for. Indeed, when assessing the impact of urbanisation on body size and colour, two variables that are measured on each individual, we are actually mixing two different questions. The first one (i.e. Model A) tests the effect of urbanisation on the bee traits at the species level (how do larger or smaller species respond to urbanisation?) whereas the second one (i.e. Model B) deals with within-species trait variation (how do larger or smaller individuals within each species respond to urbanisation?). In order to disentangle these two questions, we transformed the individual-level variables so as to obtain two different sets to include in two different versions of the same model. First, we took the average specific values of body size and the three colour variables (L*a*b*) and assigned it to each individual from a given species. Thus, all individuals from the same species had the same value for these four individual-levels variables, and we could account only for inter-specific differences in our model. In the second version of these variables, we subtracted the mean specific value of each individual variable such that the mean specific value of each species is equal to 0. This allows us to control for inter-specific variation and to account only for within-species variation in these variables.

Here, we built an initial model (model list detailed in Supp. Info. S1) with the distance to the closest natural habitat and park area having direct effects on the four individual-level variables (i.e., three colour variables and body size), on both community-level variables (i.e., species richness and abundance), and on the amount of impervious surface in a 500-m buffer. In addition, we added a direct effect of the amount of impervious surface in a 500-m buffer on all community- and individual-level variables. We also added body size as a direct predictor of the three colour components. Moreover, we specified correlated errors between our three colour variables, and between species richness and abundance. This step allows the residual errors of two variables to be correlated for a reason not explained by our model when a direct causal effect is not ecologically relevant, for example when two variables correlate with a third unknown variable. We used a linear model (LM) for the amount of impervious surface in a 500-m buffer, a generalised linear model for species richness and abundance since these variables follow a Poisson distribution, and a linear mixed-effects model (LMM) for the four individual-level variables with park ID as random intercept factor. We did not include the amount of vegetation in a 500-m buffer in our model because this variable induced a high level of collinearity in the model (VIF = 14.57). As explained above, this model was built in two versions: Model A included the mean specific values of body size and the three colour variables while Model B included the species-centred values of these same variables. This initial model therefore allows us to test the direct and indirect effects of urbanisation features on both community- and individual-level traits as well as the relationship between coloration and body size in wild bees. We then discarded the non-significant terms until we obtained the lowest values of BIC. We checked model performance using the R package *Performance* (Lüdecke et al. 2020), and we calculated marginal (fixed effect) and conditional (fixed and random effects) R^2^ for each model (Nakagawa and Schielzeth 2013).

## Results

From April to July 2016, we sampled a total of 435 wild bees belonging to 121 species, 30 genera, and 5 families, and we recorded the presence of 994 honeybees (*Apis mellifera*). We were able to successfully capture only 373 of the 435 wild bees observed. According to the Chao1 method to estimate the total species richness, observed wild bee species richness represented 52.8% of the potential maximum richness. Using Jackknife 1 and 2 indexes, the observed richness represented from 55.8% (Jackknife 2) to 68.7% (Jackknife 1) of the potential maximum richness. Finally, 55 species (45.4%) were represented by only one individual (singleton). We provide a more detailed description of bees’ ecological traits in Supp. Info. S2, and the structure of their interaction network with flowering plants in Supp. Info. S3. Our two final models resulting from the piecewise SEM approach are presented in Figure 2 and statistics are fully summarised in Supp. Info S4. Model A, including mean specific values of individual-level variables, has a Fisher’s C statistics of 26.81 with a p-value of 0.867 and 36 degrees of freedom. Similarly, Model B, including species-centred values of individual-level variables, has a Fisher’s C statistics of 22.75 with a p-value of 0.958 and 36 degrees of freedom, implying that both our models provide a good fit to our data.

**Figure 2.**
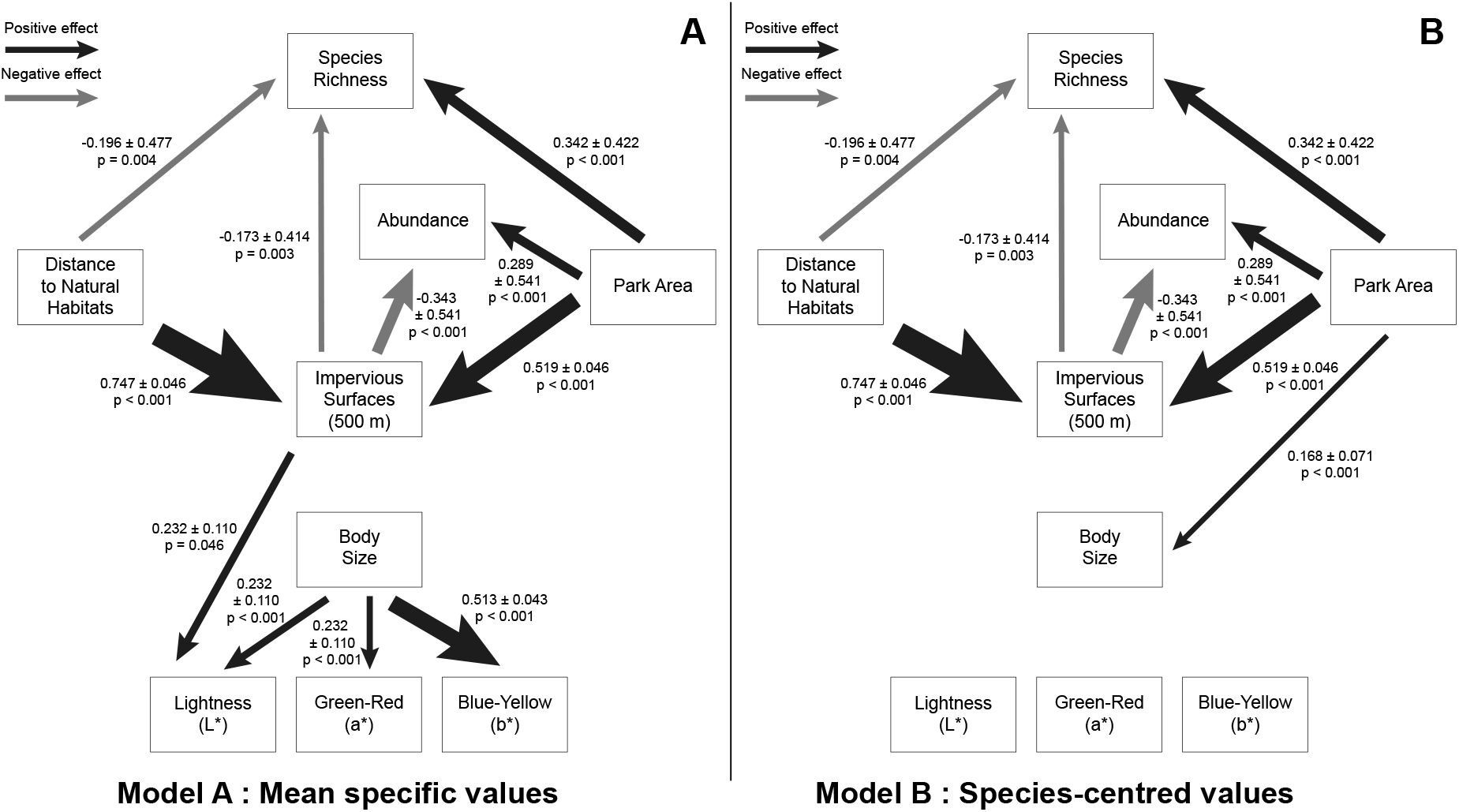
Best selected path diagrams representing the direct effects of urbanisation-related variables on the species richness, the abundance, the body size and the coloration of wild bees in the city of Marseille. Each arrow represents a statistically significant effect, which can be either negative (grey arrows) or positive (black arrows), and arrows thickness is proportional to their effect size. We provide effect size ± standard error along with the p-value. Model A (A) represents the model in which body size and the three colour variables were included as mean specific values. Model B (B) represents the same model but the values of body size and the three colour variables correspond to species-centred values (between-species variation has been eliminated by subtracting the mean specific values each time).

We found in both our models that species richness increased in larger parks (β = 0.342 ± 0.477, p = 0.004), but decreased in parks located further away from natural habitat (β = -0.196 ± 0.477, p < 0.001), and in parks surrounded by more impervious surfaces (β = -0.173 ± 0.414, p < 0.001). Abundance also increased in larger parks (β = 0.289 ± 0.541, p < 0.001) and decreased when the amount of impervious surface surrounding the park increased (β = - 0.343 ± 0.541, p < 0.001).

In addition, Model A indicated no significant relationship between average specific body size and the distance to the closest natural habitat (p = 0.075). We found that average specific lightness (L*) increased with the amount of impervious surface (β = 0.232 ± 0.110, p = 0.046). Moreover, larger bee species (average specific body size) were brighter (L*; β = 0.228 ± 0.048 p < 0.001), redder (a*; β = 0.191 ± 0.051 p = 0.001) and yellower (b*; β = 0.513 ± 0.043 p < 0.001) than smaller bee species.

In contrast, Model B showed that individuals were larger (species-centred value of body size) in larger parks (β = 0.168 ± 0.071 p = 0.028). However, we found that none of the three colour variables (species-centred values) were correlated to species-centred values of body size.

Finally, we found positively correlated errors between all three colour components for mean specific values (L* ∼ a*: β = 0.313, p < 0.001; L* ∼ b*: β = 0.774, p < 0.001; a* ∼ <b*: β = 0.708, p < 0.001), but only between L* and b* (β = 0.422, p < 0.001), and a* and b* (β = 0.341, p < 0.001) for species-centred values as L* and a* were negatively correlated (β = - 0.133, p = 0.005). We also found correlated errors between species richness and abundance (β = 0.937, p < 0.001).

## Discussion

We investigated the extent to which urbanisation impacts wild bees in the Mediterranean city of Marseille, and found that wild bees responded to urbanisation variables at the community, species, or individual level. As we detailed below, our study across biological scales provides invaluable insights into the multifaceted impacts that urbanisation has on wildlife. This integrative approach allows us to capture subtle effect variations, mainly between the inter- and intra-specific level, that would be otherwise undetectable when comparing separate studies, because of confounding factors specific to each study. We therefore encourage the use of holistic approaches across biological scales to precisely assess the impact of environmental change on animals.

The negative impact of urbanisation on bee diversity has been documented in the past (Schochet et al. 2016; Cardoso and Gonçalves 2018). Yet, some studies found no or little reduction in terms of species richness and abundance in cities (Buchholz et al. 2020; Theodorou et al. 2020). Several factors may explain these differences. For example, habitat connectivity can be highly variable among cities (Beninde et al. 2015) and may explain why some cities are more or less permeable to wildlife and bees in particular (Steffan-Dewenter and Tscharntke 1999; Buchholz et al. 2020). Furthermore, the nature and quality of the less urbanised end of the gradient can also vary. Many cities are surrounded by a more or less extended suburb and further by agricultural fields, which can have a negative impact on bee diversity depending on how crops are managed (e.g., Le Féon et al. 2010). The city of Marseille is ideal to study urbanisation gradient because it is directly surrounded by natural massifs and diversified scrubland, one of which being highly protected by the Calanques National Park, a protected area with little anthropogenic impact for bees (Ropars et al. 2020a). Therefore, we could estimate directly the extent to which wild bee assemblages penetrate into and respond to the urban matrix. More specifically, we found that the amount of impervious surface in a 500-m buffer around a park had a negative impact on both species’ richness and abundance. As impervious surfaces reduce the availability of resources and nesting sites for bees, measuring the amount of impervious surface around a park reflects its degree of isolation and its lack of connectivity with vegetation patches. This result corroborates previous findings showing that the amount of impervious surface in a 500-m buffer correlated with reduced species richness and abundance of bees (Geslin et al. 2016; Burdine and McCluney 2019b). Furthermore, our results revealed that higher species richness and abundance were found in larger city parks. This is consistent with previous studies identifying large patches of habitat as the most important factor to maintain high levels of biodiversity within cities (Beninde et al. 2015; Baldock et al. 2015; Quistberg et al. 2016). Our study thus highlights the need to create larger city parks and denser corridor networks between these parks so as to make the city of Marseille more permeable to wild bees.

Urbanisation variables did not affect the body size of bees in our study, except park size, as larger parks harboured larger individuals within species but not larger species. This effect, albeit weak based on the marginal R^2^ of the model, may reflect a higher resource availability both in terms of quality and quantity in larger parks (strong, positive correlation between park size and the amount of vegetation within parks in our data, r^2^ = 0.78), thus allowing individuals to grow larger than in smaller parks, where resources may be scarcer. This further strengthens the idea that larger parks are beneficial to bees, not only in terms of species richness and abundance, but also in terms of individual quality (Quistberg et al. 2016). With this finding, we also emphasise the need to use individual-level variables such as body size (Buchholz and Egerer 2020) to precisely assess the health of a given community of species because one can disentangle the observed effects occurring at the species level from those occurring at the individual level within species. Assessing the amount and quality of resources within parks could also improve how parks should be managed to reduce the impact of urbanisation.

In our study, we characterised the coloration of each individual we captured to assess how colour traits respond to urbanisation on one hand, and to explore the relationships between coloration and body size in bees on the other. The effect of urbanisation on animal coloration has been relatively overlooked, and although most studies focussed on birds, current evidence suggest that diurnal animals in urban areas are darker due to thermal melanism, protection against pollution, or camouflage, and display duller colour signals than their rural counterparts (e.g., Chatelain et al. 2014; Biard et al. 2017; Leveau 2021). Our results indicate that brighter (i.e., high lightness values) species are more successful than darker ones in parks surrounded by a greater amount of impervious surface. In other words, darker species are under-represented in highly urbanised areas. This is consistent with the thermal melanism hypothesis (Clusella Trullas et al. 2007) stating that darker ectotherms should be favoured in colder habitats because they heat their body up faster than bright individuals, since dark colours are more efficient at absorbing external heat. Urban environments, especially in a hot Mediterranean city such as Marseille, are particularly warm and bees are forced to live near their critical thermal maximum (Burdine and McCluney 2019a). Therefore, a possible interpretation of our results is that in the most urbanised areas, which are presumably warmer, thermoregulation is more challenging for dark species than for bright ones because they reach their critical limit too fast (Pereboom and Biesmeijer 2003). In addition, even though we cannot completely rule them out, alternative hypotheses relative to camouflage or aposematism are unlikely. First, urban-induced colour change related to camouflage usually has an opposite effect, driving urban animals towards darker coloration (Bishop and Cook 1980; Leveau 2019). Second, in the context of aposematism, having more brightly coloured species in more urbanised areas would mean that darker species are more predated in these parks. This hypothesis either implies that predation pressures in urbanised city parks is higher for darker species or lower for brighter ones compared with less urbanised city parks. Although this is plausible, this explanation is far from parsimonious and would involve too many layers of presumptions. In any case, we advocate future studies to further investigate the relationship between urbanisation and coloration in bees taking into account all ecological determinants of body coloration.

Interestingly, we found positive correlations between species body size and species coloration. More specifically, larger species are brighter, redder, and yellower while smaller species are darker, greener, and bluer. In simple terms, large species often have conspicuous colours while smaller species are much darker (Figure 3). Surprisingly perhaps, bee coloration has received relatively little attention compared with other traits but their bright coloration seems to have an aposematic function (Badejo et al. 2020). If so, our results suggest that aposematic colours are much more present in large than in small species. Two non-exclusive hypotheses could explain why larger species are more brightly coloured than smaller ones. First, aposematism signals are more efficient in large preys because predators can detect them and identify them more easily than small preys (Gamberale and Tullberg 1996 but see Remmel and Tammaru 2009). Second, larger species of bees may suffer from a higher predation pressure from birds than smaller species which are only a few millimetres in body length since birds prefer larger insect preys (Remmel and Tammaru 2009). Thus, the cost-benefit balance of producing conspicuous colours may be more advantageous for larger species than for smaller ones. Caution should be given with these possible interpretations since dark coloration can also have an aposematic function, especially when iridescent colours are involved, as demonstrated in carpenter bees from the genus *Xylocopa* (Blaimer et al. 2018).

**Figure 3.**
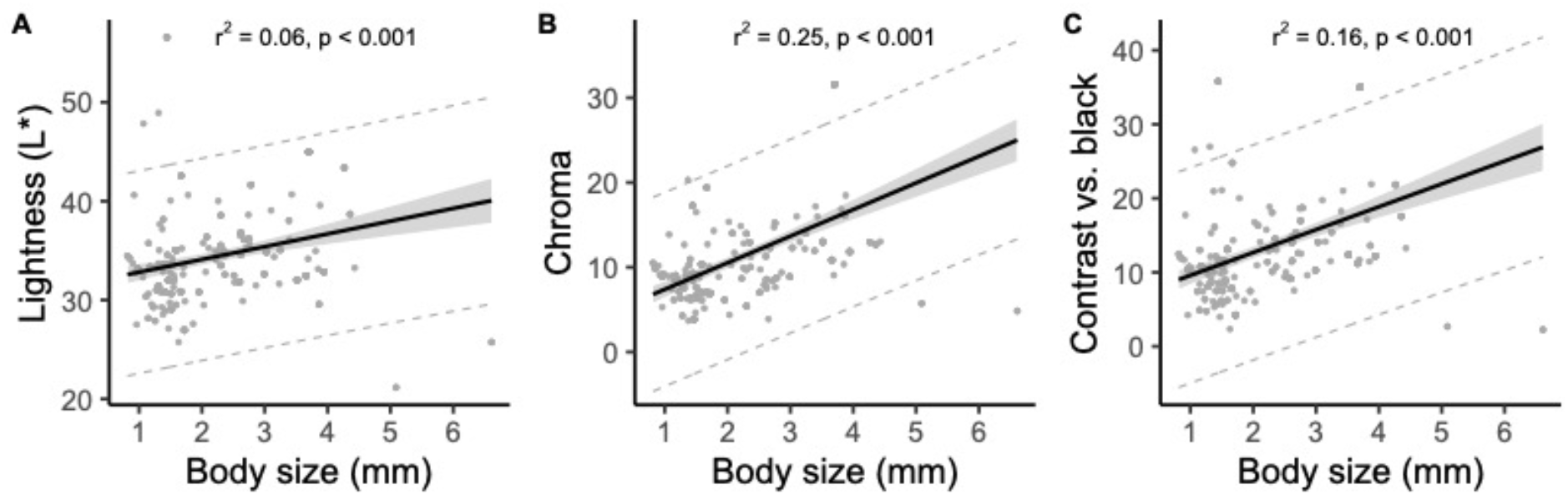
Linear regressions between colour components, i.e. lightness (A), chroma (B), and contrast against a black colour (C), and body size at the species level (mean specific values). Chroma was calculated as C = (a*^2^ + b*^2^)^1/2^ and corresponds to colour saturation. Contrasts vs. black corresponds to the distance between each colour point and a black point within the CIELAB colour space, with higher values corresponding to more colourful species. Contrast vs. black was calculated as Δ_black_ = ((L* – L*_black_)^2^ + (a* – a*_black_)^2^ + (b* – b*_black_)^2^)^1/2^. Shaded area represents the 95% confidence interval and dashed lines the 95% prediction interval. We also provide r^2^ and p-value associated with each linear regression.

To conclude, our study shows that urbanisation has a negative impact on wild bees across biological scales, with distinct responses at the community, species, and individual levels. Species richness and abundance of wild bees decrease along an urbanization in the Mediterranean city of Marseille, mainly because of the amount of impervious surface around the city parks. We also identified the size of city parks as a key factor positively affecting the wild bee community, in terms of species richness and abundance on one hand, and in the body size of individuals within species on the other. This strongly advocates for the inclusion of larger parks in city centres to maintain acceptable levels of biodiversity. Brighter species are also more successful in urbanised areas, perhaps due to the thermal advantage that their bright colours confer them, suggesting that coloration is an important trait to consider when assessing the impact of environmental change of functional diversity. Finally, we uncovered a positive correlation between species size and colour in wild bees and urge future studies to explore the details and ecological function of these relationships.

## Acknowledgements

We thank the gardeners and the stakeholders of the urban parks of Marseille for helping us during fieldwork. We particularly appreciated the help of Patrick Bayle, Catherine Stenou and Josette Sakakini. We thank David Genoud, Eric Dufrêne and Matthieu Aubert for species identification. We are also grateful to Léa Chalvin & Rémy Roques for their help during fieldwork.

## Declarations

### Funding

The authors have no funding to acknowledge for this study

### Conflict of interests

The authors have no conflict of interest to declare that are relevant to the content of this article

### Authors’ contributions

A.B., L.R., L.S., F.F. and B.G. conceived the study. L.S., M.D.C., C.R., M.Z. and B.G. participated in fieldwork or data collection. A.B. and L.R. performed the statistical analyses and L.R. extracted landscape variables. A.B. wrote the manuscript with L.R., F.F. and B.G. and all authors reviewed it and provided feedback.

### Consent for publication

All the authors consent for the publication of this manuscript

## Notes

### Competing Interest Statement

The authors have declared no competing interest.

