## Supplementary information for "Urbanisation impacts the diversity, coloration, and body size of wild bees in a Mediterranean city"

**Supp. Info. S1**

List of linear mixed models included in the initial piecewise SEM procedure. We indicate the modelling method, the response variables, the predictors and the random intercept used in each model. This model was built in two slightly different versions. In the first one, we included body size and the three colour variables as average specific values, and in the second one we included the within-species variation values. LM = linear model; GLM = generalised linear model; LMM = linear mixed-effects model.

| Method | Response | Predictors | Random |
|---|---|---|---|
| LM | Impervious surfaces (500 m) | Distance-Wild + Park area | - |
| GLM | Species Richness | Distance-Wild + Park area + Impervious-500m | - |
| GLM | Abundance | Distance-Wild + Park area + Impervious-500m | - |
| LMM | Body Size | Distance-Wild + Park area + Impervious-500m | Park ID |
| LMM | Lightness (L*) | Distance-Wild + Park area + Impervious-500m + Body size | Park ID |
| LMM | Green-Red (a*) | Distance-Wild + Park area + Impervious-500m + Body size | Park ID |
| LMM | Blue-Yellow (b*) | Distance-Wild + Park area + Impervious-500m + Body size | Park ID |

**Supp. Info. S2**

Richness estimation of bee species

| Variables | Value | 95% confidence interval |
|---|---|---|
| **Bee abundance** | 1429 | - |
| **Number of bee species** | 121 | - |
| **Chao1** | 229 | [172; 347] |
| **Jackknife1** | 176 | [159; 201] |
| **Jackknife2** | 217 | [187; 260] |

Excluding honey bees, the *Lasioglossum* genus dominated the sample representing 19.8% of wild bee abundance and being the most diverse genus with 18 species. The most common species was *Bombus pascuorum* with 29 individuals, corresponding to 6.7% of the total wild bees captured followed by *Xylocopa violacea* with 22 individuals (5.0%).

Considering the ecological traits of bee species, we recorded 50 belowground nesting species, 49 aboveground nesting species and 8 kleptoparasitic species corresponding to 41.3%, 40.4% and 6.6% of the total bee species richness. Nesting habits was unknown for 15 species (12,4%). Considering the floral preferences, 79 species were generalist and foraged on a wide range of plant species (65.3%) and 21 species were specialist and foraged on one or few plant species (17.3%). The floral diet is not known for 22 species (18.2%).

A European IUCN status was available for 106 species (excluding species complexes and unidentified species). Our study included 84 least concerned species (LC – 79.2%), 21 data deficient species (DD – 18.8%), and 3 near-threatened species (NT – 2.8%; *Andrena ovatula*, *Lasioglossum minutulum* and *Lasioglossum prasinum*). Finally, 5 species over the 121 were endemic to Europe (*Ceratina gravidula*, *Halictus crenicornis*, *Hylaeus cf. pictus*, *Lasioglossum minutulum* and *Panurgus dentipes*).

**Supp. Info. S3 :**

Interaction network between plant species (left part) and bee species (right part) within the city of Marseille. Threatened pollinator species and their links are highlighted in red, while european endemic bee species are highlighted in blue. Plant highlighted in green corresponds to cultivated or non native species. To characterize this plant-bee network, we calculated modularity, nestedness, number of compartment and connectance using the *computeModules* and *networklevel* functions from the *bipartite* package (Dormann et al. 2009). The most generalist and dominant bee species, *Apis mellifera*, is highlighted in yellow. The plant-bee network includes the 121 bee species in interaction with 103 plant species representing 360 links. The network has 7 compartments with six pairs of plant-bee interactions which are isolated from the rest of the network. The modularity, nestedness and connectance of the network were estimated at 0.315, 1.626 and 0.028, respectively. The network is highly dominated by *Apis mellifera*, which forages on 60 plant species. On the other hand, *Bellis perennis*, *Trifolium repens*, *Malva sylvestris* and *Echium vulgare* appeared as central nodes within the network, supporting respectively 23, 22, 16 and 14 wild bee species.

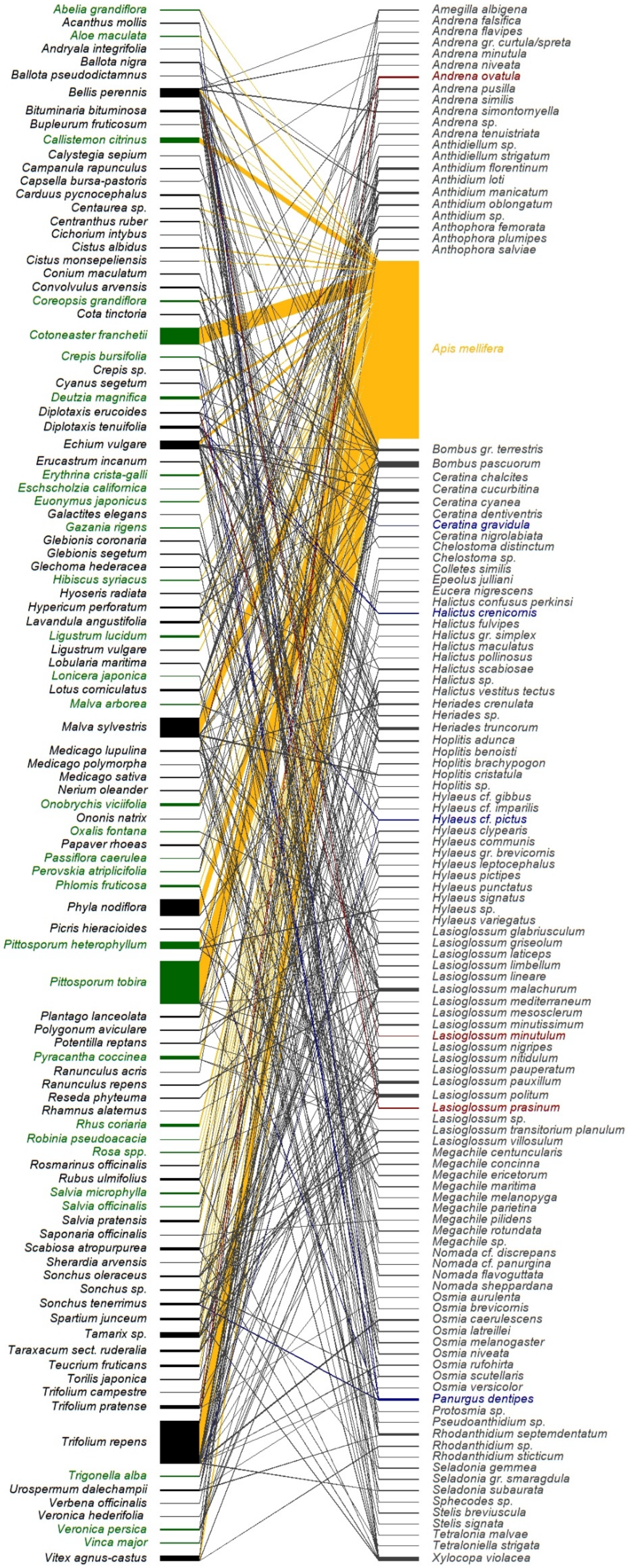
Abelia grandiflora
Acanthus mollis
Aloe maculata
Andryala integrifolia
Ballota nigra
Ballota pseudodictamnus
Bellis perennis
Bituminaria bituminosa
Bupleurum fruticosum
Callistemon citrinus
Calystegia sepium
Campanula rapunculus
Capsella bursa-pastoris
Carduus pycnocephalus
Centaurea sp.
Centranthus ruber
Cichorium intybus
Cistus albidus
Cistus monspeliensis
Conium maculatum
Convolvulus arvensis
Coreopsis grandiflora
Cota tinctoria
Cotoneaster franchetii
Crepis bursifolia
Crepis sp.
Cyanus segetum
Deutzia magnifica
Diplotaxis erucoides
Diplotaxis tenuifolia
Echium vulgare
Erucastrum incanum
Erythrina crista-galli
Eschscholzia californica
Euonymus japonicus
Galactites elegans
Gazania rigens
Glebionis coronaria
Glebionis segetum
Glechoma hederacea
Hibiscus syriacus
Hyoseris radiata
Hypericum perforatum
Lavandula angustifolia
Ligustrum lucidum
Ligustrum vulgare
Lobularia maritima
Lonicera japonica
Lotus corniculatus
Malva arborea
Malva sylvestris
Medicago lupulina
Medicago polymorpha
Medicago sativa
Nerium oleander
Onobrychis viciifolia
Ononis natrix
Oxalis fontana
Papaver rhoeas
Passiflora caerulea
Perovskia atriplicifolia
Phlomis fruticosa
Phyla nodiflora
Picris hieracioides
Pittosporum heterophyllum
Pittosporum tobira
Plantago lanceolata
Polygonum aviculare
Potentilla reptans
Pyracantha coccinea
Ranunculus acris
Ranunculus repens
Reseda phyteuma
Rhamnus alaternus
Rhus coriaria
Robinia pseudoacacia
Rosa spp.
Rosmarinus officinalis
Rubus ulmifolius
Salvia microphylla
Salvia officinalis
Salvia pratensis
Saponaria officinalis
Scabiosa atropurpurea
Sherardia arvensis
Sonchus oleraceus
Sonchus sp.
Sonchus tenerrimus
Spartium junceum
Tamarix sp.
Taraxacum sect. ruderalia
Teucrium fruticans
Torilis japonica
Trifolium campestre
Trifolium pratense
Trifolium repens
Trigonella alba
Urospermum dalechampii
Verbena officinalis
Veronica hederifolia
Veronica persica
Vinca major
Vitex agnus-castus
Amegilla albigena
Andrena falsifica
Andrena flavipes
Andrena gr. curtula/spreta
Andrena minutula
Andrena niveata
Andrena ovatula
Andrena pusilla
Andrena similis
Andrena simontornyella
Andrena sp.
Andrena tenuistriata
Anthidiellum sp.
Anthidiellum strigatum
Anthidium florentinum
Anthidium loti
Anthidium manicatum
Anthidium oblongatum
Anthidium sp.
Anthophora femorata
Anthophora plumipes
Anthophora salviae
Apis mellifera
Bombus gr. terrestris
Bombus pascuorum
Ceratina chalcites
Ceratina cucurbitina
Ceratina cyanea
Ceratina dentiventris
Ceratina gravidula
Ceratina nigrolabiata
Chelostoma distinctum
Chelostoma sp.
Colletes similis
Epeolus julliani
Eucera nigrescens
Halictus confusus perkinsi
Halictus crenicornis
Halictus fulvipes
Halictus gr. simplex
Halictus maculatus
Halictus pollinosus
Halictus scabiosae
Halictus sp.
Halictus vestitus tectus
Heriades crenulata
Heriades sp.
Heriades truncorum
Hoplitis adunca
Hoplitis benoisti
Hoplitis brachypogon
Hoplitis cristatula
Hoplitis sp.
Hylaeus cf. gibbus
Hylaeus cf. imparilis
Hylaeus cf. pictus
Hylaeus clypearis
Hylaeus communis
Hylaeus gr. brevicornis
Hylaeus leptocephalus
Hylaeus pictipes
Hylaeus punctatus
Hylaeus signatus
Hylaeus sp.
Hylaeus variegatus
Lasioglossum glabriusculum
Lasioglossum griseolum
Lasioglossum laticeps
Lasioglossum limbellum
Lasioglossum lineare
Lasioglossum malachurum
Lasioglossum mediterraneum
Lasioglossum mesosclerum
Lasioglossum minutissimum
Lasioglossum minutulum
Lasioglossum nigripes
Lasioglossum nitidulum
Lasioglossum pauperatum
Lasioglossum pauxillum
Lasioglossum politum
Lasioglossum prasinum
Lasioglossum sp.
Lasioglossum transitorium planulum
Lasioglossum villosulum
Megachile centuncularis
Megachile concinna
Megachile ericetorum
Megachile maritima
Megachile melanopyga
Megachile parietina
Megachile pilidens
Megachile rotundata
Megachile sp.
Nomada cf. discrepans
Nomada cf. panurgina
Nomada flavoguttata
Nomada sheppardana
Osmia aurulenta
Osmia brevicornis
Osmia caerulescens
Osmia latreillei
Osmia melanogaster
Osmia niveata
Osmia rufohirta
Osmia scutellaris
Osmia versicolor
Panurgus dentipes
Protosmia sp.
Pseudoanthidium sp.
Rhodanthidium septemdentatum
Rhodanthidium sp.
Rhodanthidium sticticum
Seladonia gemmea
Seladonia gr. smaragdula
Seladonia subaurata
Sphecodes sp.
Stelis breviuscula
Stelis signata
Tetralonia malvae
Tetraloniella strigata
Xylocopa violacea

**Supp. Info. S4**

Table providing the full results from our final model resulting from the piecewise SEM procedure. For each test, we provide the name of the response and predictor variables, the standardised $\beta$ estimates and their associated standard errors (SE), the p-value, and the marginal ($R^2m$) and conditional ($R^2c$) $R^2$. Averaged specific values of each individual variables are indicated with (sp) while within-species variation values are indicated as (ind). (sp) and (ind) values were run in two separate models otherwise identical. Significant p-values are indicated in bold.

| Response | Predictor | $\beta$ ± SE | *p* | $R^2$m | $R^2$c |
|---|---|---|---|---|---|
| Impervious surfaces | Distance-Wild | 0.747 ± 0.046 | **< 0.001** | 0.43 | - |
| | Park area | 0.519 ± 0.046 | **< 0.001** | | |
| Species Richness | Distance-Wild | -0.196 ± 0.477 | **0.004** | 1 | - |
| | Park area | 0.342 ± 0.422 | **< 0.001** | | |
| | Impervious surfaces | -0.173 ± 0.414 | **0.003** | | |
| Abundance | Park area | 0.289 ± 0.541 | **< 0.001** | 1 | - |
| | Impervious surfaces | -0.343 ± 0.541 | **< 0.001** | | |
| Body Size (sp) | Distance-Wild | 0.753 ± 0.073 | 0.075 | 0.01 | 0.05 |
| Lightness (L*) (sp) | Impervious surfaces | 0.232 ± 0.110 | **0.046** | 0.11 | 0.25 |
| | Body Size (sp) | 0.228 ± 0.048 | **< 0.001** | | |
| Green-Red (a*) (sp) | Body Size (sp) | 0.191 ± 0.051 | **< 0.001** | 0.04 | 0.05 |
| Blue-Yellow (b*) (sp) | Impervious surfaces | 0.137 ± 0.079 | 0.138 | 0.29 | 0.35 |
| | Body Size (sp) | 0.513 ± 0.043 | **< 0.001** | | |
| Body Size (ind) | Distance-Wild | 0.107 ± 0.072 | 0.158 | 0.02 | 0.04 |
| | Park area | 0.168 ± 0.071 | **0.028** | | |
| Lightness (L*) (ind) | Park area | -0.081 ± 0.052 | 0.132 | 0.01 | 0.01 |
| Green-Red (a*) | Impervious surfaces | -0.073 ± 0.66 | 0.280 | 0.01 | 0.03 |

| | | | | | |
|---|---|---|---|---|---|
| (ind) | | | | | |
| Blue-Yellow (b*) (ind) | Body Size (ind) | 0.058 ± 0.052 | 0.261 | 0.00 | 0.04 |
